# A natural soil-derived microbiota reshapes nitrogen form and plant mineral nutrition

**DOI:** 10.64898/2026.05.07.723502

**Authors:** Laura Dethier, Jiaen Xie, Xiaoran Yang, Yang Bai, Deyang Xu

## Abstract

Nitrogen form in soil has a major influence on plant growth and mineral nutrition, yet whether and how nitrogen form is controlled by microbiota in natural soils remains poorly understood. Here we show that, in a high-organic-matter soil from Danish nature, the native soil microbiota determines nitrogen form and thereby controls plant mineral nutrition. Eliminating the microbiota by sterilization disrupted nitrification, causing ammonium accumulation and loss of nitrate, which resulted in impaired growth and a pronounced reduction in shoot Mg and Ca associated with chlorosis. Reintroduction of a simplified soil-derived microbiota restored nitrification and re-established a balanced NO□□/NH□□ regime, which rescued Mg and Ca accumulation, alleviated chlorosis, and promoted plant growth. Metagenomic analyses of bulk soil, rhizosphere, and root-associated communities revealed enrichment of nitrogen-cycling functions, including nitrification-related genes, supporting the capacity of the restored microbiota to modulate nitrogen form in soil and the rhizosphere. Moreover, this microbiota alleviated mineral deficiency symptoms in an organic agricultural soil. Together, our findings reveal a natural microbiota-dependent mechanism by which soil microbes determine nitrogen form and thereby regulate plant mineral nutrition, particularly Mg and Ca homeostasis.

## Main text

Nitrogen (N) is a major macronutrient required for plant growth and a central component of proteins, nucleic acids and chlorophyll(Hawkesford et al. 2012; de Bang et al. 2021). Plants predominantly take up inorganic N in the form of nitrate (NO_3_^-^) and ammonium (NH□□), but most species do not grow optimally on NH□□ as the sole N source(Roosta and Schjoerring 2007; Roosta et al. 2009). High NH□□ levels in growing medium can compete with cations, such as K^+^, Mg^2+^ and Ca^2+^, thus reducing their uptake(Roosta and Schjoerring 2007; Hernández-Gómez et al. 2015; Coleto et al. 2023). In contrast, NO_3_^-^ or mixed N forms generally facilitate plant growth under high N supply, and also enhance the uptake of essential cations(Roosta and Schjoerring 2007; Roosta et al. 2009). Therefore, not only the quantity but also the form of nitrogen in soil and their availability to roots are key for plant growth and nutrition(Roosta and Schjoerring 2007). Microbes play a central role in shaping plant N acquisition(Zhang et al. 2019; Li et al. 2026; Ma et al. 2026). In soils, the balance between NO_3_^-^ and NH□□ is largely determined by microbial processes such as mineralization of organic N, nitrification, denitrification and N fixation(Sieradzki et al. 2023; Li et al. 2024; Ma et al. 2026). However, how the microbiota regulates nitrogen form and thereby controls both plant growth and mineral nutrition remains underexplored. Here, we show that in the absence of a nitrifying microbiota, ammonium accumulates in the soil, leading to poor plant growth and reduced Mg and Ca accumulation in shoots. Reintroduction of a simplified microbial community via soil suspension restored nitrification and a more balanced NO□^-^/NH□□ regime, thereby improving plant biomass and alleviating Mg and Ca deficiency-associated phenotypes.

To examine how soil microbiota influence plant mineral nutrition, we grew *Arabidopsis* in natural high-organic-matter soil (75% organic matter) collected from an alder swamp in North Zealand, Denmark, either sterilized by gamma irradiation (S) or left untreated (NS), and compared plant performance between treatments. Plants grown in S soil developed chlorosis on the margins of older leaves (Fig. 1a), indicative of impaired nutrient homeostasis. We therefore quantified soil nitrate-N (NO□^-^_-_N), ammonium-N (NH□□-N), Olsen phosphorus (inorganic P) and exchangeable cations (Mg, Ca and K) at the start of the experiment and after five weeks of plant growth. Sterilized soil contained more inorganic N than NS soil at both time points (P < 0.05; Fig. 1c). In S soil, inorganic N was predominantly present as NH□□-N and remained unchanged between week 0 and week 5 (Wilcoxon rank-sum test, P > 0.05), whereas NS soil contained only NO□^-^-N, consistent with the disruption of microbial N transformations by sterilization. Notably, inorganic N declined over time in NS soil (P < 0.01; Fig. 1c, Table S1). Inorganic P was also higher in S soil at both sampling points (P < 0.01), whereas exchangeable cation concentrations were comparable at week 0 but lower in NS soil after five weeks (P < 0.05; Table S1). Together, these results indicate that, in this soil with high organic matter, the native microbiota is required for N transformation.

**Fig. 1.**
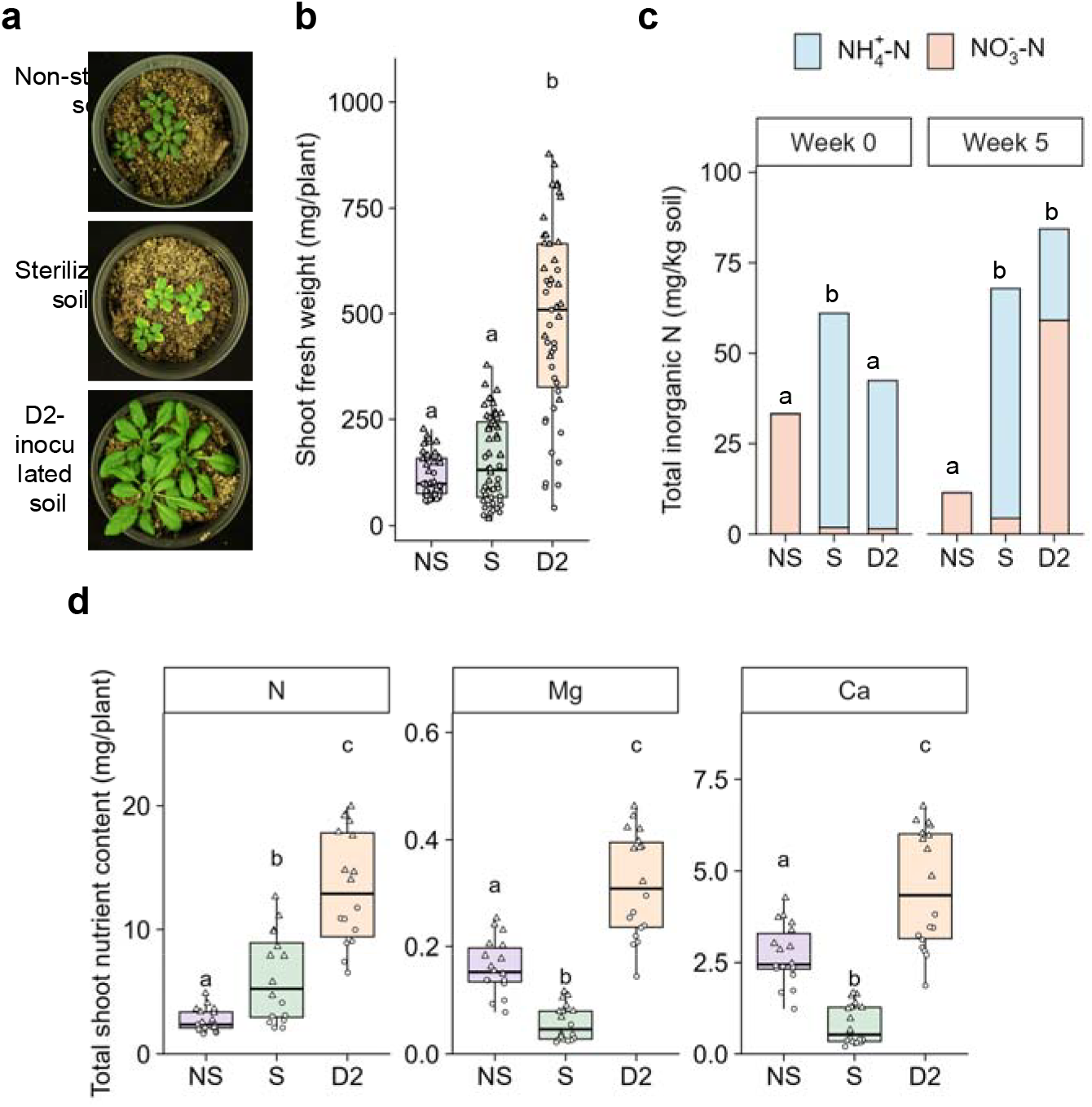
A soil-derived microbiota enhances *Arabidopsis* growth, nutrient accumulation, and soil nitrification. **a**, Representative images of *Arabidopsis* grown in non-sterilized (NS) soil, sterilized (S) soil, or S soil inoculated with a soil suspension (D2) for 5 weeks. **b**, Shoot fresh weight of five-week-old plants grown under the indicated treatments (n = 54, 43 and 47 plants from left to right). **c**, Total inorganic nitrogen (NO_3_^-^-N and NH_4_^+^-N) concentrations in bulk soil at week 0 and week 5. Data represent the mean (n = 6 per treatment). **d**, Total shoot nitrogen (N), magnesium (Mg), and calcium (Ca) content measured by ICP-OES (n = 18 per treatment). Data are pooled from two independent experiments, represented by different shapes in **b** and **d**. Different letters indicate significant differences (P < 0.05; Kruskal-Wallis test followed by Dunn’s post hoc test with Benjamini-Hochberg correction).

Leaf elemental analysis revealed contrasting shoot nutrient profiles between treatments. Both the concentrations and total amounts of N, P and K were higher in the rosette leaves of plants grown in S soil than in those grown in NS soil (P < 0.01, Fig. 1d, Fig. S1). Leaf concentrations of Fe, Mn, Mo, and S were also higher in S plants (P < 0.05, Fig. □S2). Notably, plants grown in S soil showed reduced Mg (∼210%) and Ca (∼302%) concentrations, resulting in a lower total Mg and Ca per plant (P < 0.001, Fig.1d, Fig. S1). This decrease in Mg is consistent with the observed chlorosis phenotype (Fig.1a), as Mg deficiency typically appears first in older leaves due to the high phloem mobility of Mg(de Bang et al. 2021). Supplying plants with high NH□^+^ has been shown to reduce leaf concentrations of inorganic cations, including Ca and Mg, while increasing the accumulation of P and S anions in order to maintain charge balance(Roosta and Schjoerring 2007; Roosta et al. 2009; Hernández-Gómez et al. 2015). In line with this, the shoot nutrient profile of plants grown in S soil is consistent with the high NH□^+^ levels measured in the soil. Together, these results suggest that sterilization inhibits nitrification in the soil, leading to NH□□ accumulation, which might cause the reduction of Mg and Ca uptake and chlorosis.

We next inoculated S soil with serial dilutions of NS soil to test how decreasing microbial diversity from NS soil affects plant growth. This soil dilution approach has been used in several studies to assess how microbial diversity loss impacts soil functions, including nitrification and denitrification(Griffiths et al. 2001; Philippot et al. 2013; Chen et al. 2020; Yang et al. 2023). We found that inoculating S soil with the original suspension (D1) reduced chlorosis and increased shoot fresh weight by ∼534% relative to plants grown in S soil without inocula (P < 0.001), whereas higher dilution treatments (10^-2^-to 10^-10^-fold dilutions of D1) had no significant effects on leaf color and biomass (Fig. S3). In a separate experiment, diluting D1 twofold (D2) also rescued chlorosis and promoted shoot growth relative to control suspensions prepared from sterilized or heat-treated soil (P < 0.001), while higher dilutions of D1 (4- to 64-fold dilutions) had no effects on plant phenotype (Fig. S3).

Thus, these results identify D2 as the most diluted microbial inoculum that rescues chlorosis and promotes plant growth.

Balanced nitrogen forms (i.e. NO_3_^-^ and NH□□) can facilitate plant growth compared to a single N source(Roosta and Schjoerring 2007). We therefore hypothesized that the chlorosis rescue and growth promotion observed in D2 relative to S and NS soil (Fig. 1a-b) are driven by balancing nitrogen form. We found soil nutrient levels to be similar between D2 and S soil at week 0 (P > 0.05, Table S1), confirming that the inoculum did not introduce nutrients to the soil. After five weeks, NO_3_^-^-N levels increased 41-fold in D2 compared to week 0 (Wilcoxon rank-sum test, p < 0.001), while NH□ □-N levels slightly decreased (P > 0.05, Fig. 1c, Table S1). Available P decreased slightly over the same period (P < 0.05), while exchangeable cations remained stable and did not differ from S soil at the experimental end point (P > 0.05, Table S1). Rosette elemental profiling showed that D2 plants contained higher concentrations of Mg (∼88%) and Ca (∼103%) than S plants, as well as higher N (∼53%), P (∼128%), and K (∼117%) concentrations than NS plants (P < 0.001, Fig. S1). Furthermore, total amount of Mg, Ca, N, P and K was the highest in D2 plants (P < 0.05, Fig. 1d, Fig. S1). As the exchangeable soil cation pools were unchanged between S and D2 soils (Table S1), the increased Mg and Ca content in D2 plants suggests enhanced uptake of these cations driven by microbial transformation of NHL^+^ to NO_3_ ^-^ in soil. Moreover, a balanced N supply, rather than either NO_3_ ^-^ or NH□^+^ alone, promotes optimal plant growth.

To test whether the high soil nitrate levels observed in D2 treatment could explain the PGP effect and chlorosis rescue, S soil was supplemented with increasing concentrations of KNO_3_ (0, 60, 120, and 240 mg N kg^-1^ soil) as a nitrate source. In parallel, S soil was amended with Mg, Ca or a combination of both. All KNO_3_ treatments rescued chlorosis and increased shoot fresh weight by ∼297%, ∼284%, and ∼348%, respectively, compared to plants grown without nitrate (P < 0.05, Fig. S4), indicating that nitrate availability is sufficient to reproduce the D2 phenotype. Additionally, soil pH in both nitrate-treated and untreated soils remained between 6.0 and 6.5. This range does not limit nor substantially affect Mg and Ca availability(Gransee and Führs 2013), indicating that pH changes did not contribute to the observed rescue effect. In contrast, supplementation with Mg, Ca, or their combination did not rescue chlorosis nor increase shoot biomass relative to the control (Fig. S4). Together, these results suggest that chlorosis in plants grown in S soil results from nitrogen imbalance rather than from a deficiency of Ca and Mg, and that restoration of nitrate availability is sufficient to restore plant growth and nutrient status.

We next investigated the microbiota composition and their function using 16S rRNA gene amplicon sequencing and metagenomic sequencing of soils and five-week-old plants grown in S, NS and D2 soils. Beta diversity analysis revealed that community composition was primarily driven by treatment (R^2^ = 0.395, F = 45.95, P = 0.001), with a smaller but significant effect of experiment (R^2^ = 0.086, F= 19.98, P = 0.001, Table S3). Pairwise PERMANOVA confirmed that all treatments differed significantly from each other, and this pattern was consistent across the two independent experiments (P = 0.001). This separation was reflected in the PCoA, particularly for NS samples, which were clearly distinct from D2 and S (Fig. 2a). Community composition differed between root and rhizosphere (R^2^ = 0.077, F = 8.74, P = 0.0015), and between root and bulk soil (R^2^ = 0.074, F = 5.56, P = 0.0015), but not between rhizosphere and bulk soil compartment (R^2^ = 0.016, F = 1.14, P = 0.291, Table S3). Treatment significantly explained variation within all three compartments (P = 0.001). Alpha diversity was highest in NS treatment for all compartments (Fig. 2b, Fig. S5). D2 partially restored richness and Chao1 in rhizosphere and roots relative to S (P < 0.05, Fig. 2b), while Shannon evenness did not differ between S and D2 (P > 0.05, Fig. S5). Taxonomic composition at the phylum level was dominated by *Pseudomonadota, Actinomycetota, Bacillota* and *Bacteroidota* in all treatments, however, D2 showed an increased relative abundance of *Bacillota*, particularly in the rhizosphere (Fig. 2c, Table S4). Together, these results indicate that D2 establishes a simplified but distinct microbiota with reduced diversity compared to the donor NS soil.

**Fig. 2.**
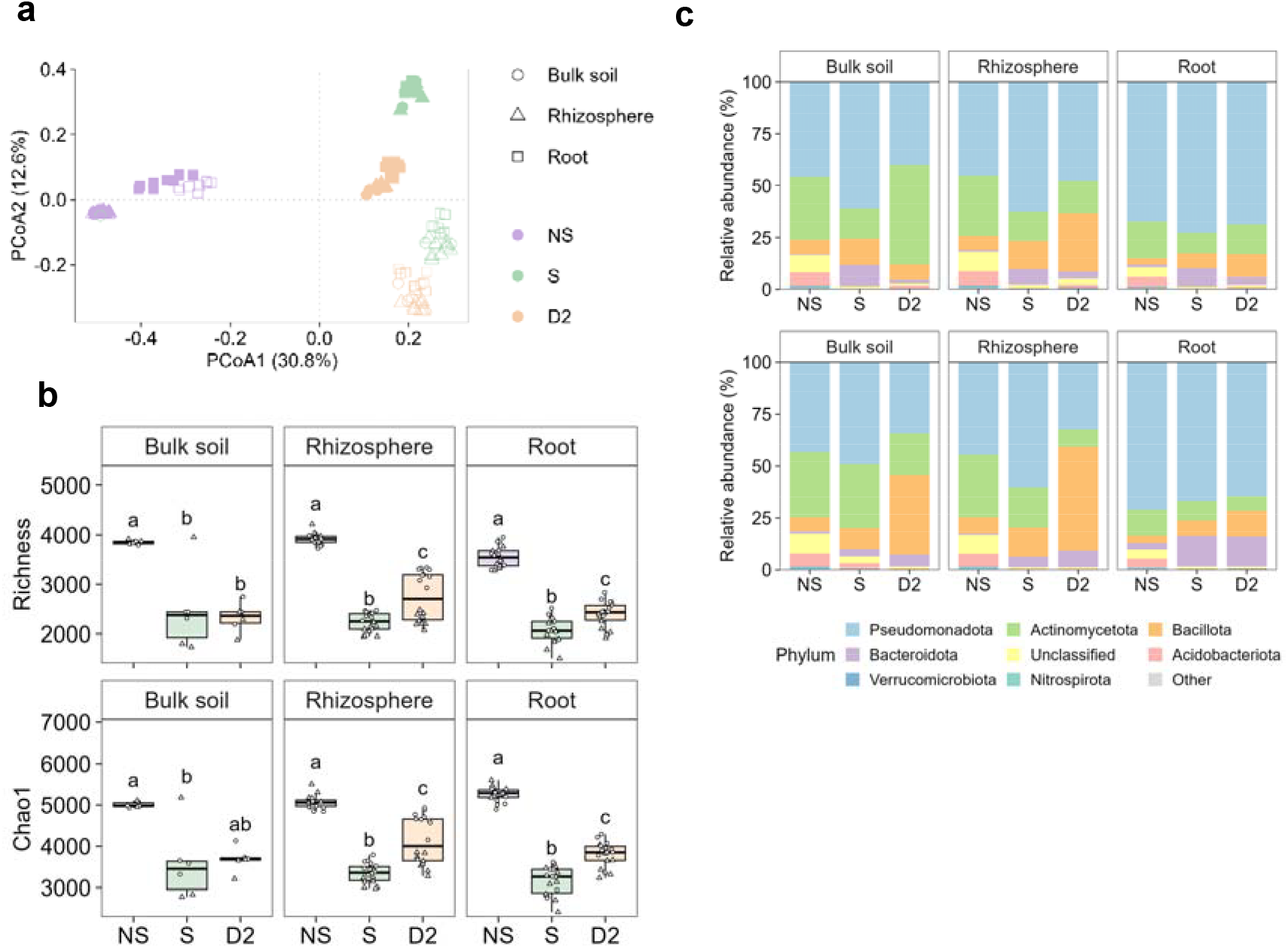
A soil-derived microbiota establishes a distinct bacterial community across soil and plant compartments. **a**, Principal coordinate analysis (PCoA) based on Bray-Curtis dissimilarities showing bacterial community composition in bulk soil, rhizosphere, and root compartments of *Arabidopsis* grown under NS, S, or D2 conditions for five weeks. **b**, Alpha diversity measured as richness (top) and Chao1 index (bottom) across compartments and treatments. **c**, Relative abundance of bacterial taxa at the phylum level. The seven most abundant phyla are shown; remaining taxa are grouped as “Unclassified” and “Other”. Data are pooled from two independent experiments (n = 6 for bulk soil, n = 17-18 for rhizosphere, and n = 18 for root per treatment). In **a**, experiments are indicated by filled and hollow shapes; in **b**, experiments are indicated by different shapes; in **c**, the top and bottom panels correspond to experiment 1 and experiment 2, respectively. In **b**, different letters show statistically significant differences (P < 0.05; Kruskal-Wallis test with Dunn’s post hoc test and Benjamini-Hochberg correction).

Despite differences in community composition across experiments, D2 consistently promoted plant growth and rescued chlorosis. To assess the functional potential of D2 on soil N form, we curated a collection of genes involved in the cycling of inorganic N, grouped into 6 pathways, i.e. N fixation, nitrification, denitrification, dissimilatory nitrate reduction (DNR), and assimilatory nitrate reduction (ANR), and anammox(Nelson et al. 2015; Tu et al. 2019; Li et al. 2024; Dai et al. 2025) (Table S5). As the alder swamp soil is high in organic matter, we also collected genes involved in the degradation of high molecular weight organic N compounds into amino acids and oligopeptides, and their subsequent mineralization into NH□^+^ (Noll et al. 2019) as two additional pathways. These genes encode extracellular enzymes, such as proteases, peptidases, ureases, chitinases, nucleases and glutaminase(Ricketts et al. 2016; Sieradzki et al. 2023) (Table S5). Metagenomic reads were annotated against the KEGG database, and differential abundance of N-cycling KEGG orthologs (KOs) was assessed.

We compared the relative abundance of KOs within individual pathways and at the whole pathway level (defined as the sum of individual KOs’ relative abundance) in all compartments. In bulk soil, D2 showed the highest relative abundance of the organic N degradation pathway (Fig. S6). However, none of the KOs showed significant differences in their relative abundance between D2 and S, although the four nitrification genes (amoA, *amoB, amoC, hao*) were detected in all soil treatments (Table S6). In contrast, NS showed an enrichment of 14 KOs relative to S, including *amoB*, and *amoC* (log2FC: 1.41-1.51, FDR < 0.05). In the rhizosphere, 13 KOs were enriched in D2 compared to S, including *amoB* and *hao* (log2FC: 1.48-1.84, FDR < 0.001, Table S6). NS showed 23 enriched KOs, including all four nitrification genes (log2FC: 1.75 to 2.15, FDR < 0.001), of which 10 were shared with D2. Notably, three organic N degradation genes encoding chitinases and lysozymes (chitinase [EC:3.2.1.14], *acm, chiA*) were uniquely enriched in D2. In roots, 20 KOs were enriched in D2 relative to S, including *amoB* and *hao* (log2FC: 0.88-1.02, FDR < 0.05). NS showed 25 enriched KOs, 17 of which were shared with D2 and involved in denitrification, DNR, nitrification, and N fixation. Consistent with the rhizosphere, *acm* and chitinase [EC:3.2.1.14] were uniquely enriched in D2, alongside *narB* involved in ANR. Thus, the enrichment of N cycling KOs in D2 root and rhizosphere supports that the D2-derived microbiota has the functional capacity to modulate N form and contribute to nitrification of highly abundant NH□□ in the rhizosphere and soil environment.

Finally, to assess whether the beneficial effect of D2 extends to other soil types, we inoculated an autoclaved and non-autoclaved organic agricultural soil with D2 suspensions and further dilution series (equivalent to five and tenfold dilutions of D1). *Arabidopsis* grown in autoclaved soil without inoculum developed chlorosis on the leaf margins, which was rescued by D2 inoculation, while chlorosis was absent in non-autoclaved soil irrespective of the inoculum (Fig. S7). In autoclaved soil, all suspensions increased shoot biomass relative to the control (P < 0.01), while no growth promotion was observed in non-autoclaved soil (P > 0.05). This suggests that the inhibition of nitrifiers through soil sterilization is common to different soil types and that the nitrifier population from D2 can restore growth and rescue chlorosis in other soils.

In agriculture, more than half of applied N is lost through leaching and gaseous losses including ammonia volatilization and nitrous oxide emissions(Coleto et al. 2023). Nitrate is more prone to leaching than ammonium as it is not adsorbed onto soil particles and therefore remains in the soil solution(Hawkesford et al. 2012; Qiao et al. 2015). Thus, strategies that inhibit microbial nitrification to retain more ammonium in the soil have been proposed to reduce N losses but complete inhibition of nitrification is also not desirable(Qiao et al. 2015; Subbarao and Searchinger 2021; Kuppe and Postma 2024). Our study shows that the D2 community obtained through soil suspensions can enhance nitrification in soil, which improved plant growth and promoted the uptake of Mg and Ca. Interestingly, plant growth promotion and chlorosis rescue were already lost at twofold dilution of D2 (≥ 4-fold of stock suspension), suggesting that key taxa required for nitrification are sensitive to dilution and that this function is not fully redundant in this soil. Contrasting effects of microbial diversity loss on soil N cycling were observed in previous studies, including cases were strong reductions in species richness had no impact on denitrification and nitrification(Wertz et al. 2006; Chen et al. 2020) as well as cases where highly diluted communities showed reduced denitrification and nitrification(Griffiths et al. 2001; Philippot et al. 2013). This could be due to differences in soil type, pH, nutrient conditions, biotic interactions and the initial community composition(Philippot et al. 2013; Chen et al. 2020; Yang et al. 2023). Our results highlight the importance of maintaining nitrification for balanced nutrient uptake, especially of Mg and Ca, and for promoting plant growth, and provide insight for future development of soil-derived, microbiota-mediated strategies to improve plant growth and nutrition in the green transition.

## Materials and Methods

### Soil collection, processing, and sterilization

Soil was collected in March 2020 from an alder swamp in North Zealand, Denmark (55°58′05.6″ N, 12°16′16.7″ E). The soil is characterized by a 75% organic matter content, 22% sand, 1% clay, no silt, a pH of 5.4, and a cation exchange capacity (CEC) of 100 cmol_c_ kg ^-1^. Sampling, processing, and γ-irradiation was described previously(Dethier et al. 2025). Briefly, the upper 0-30 cm of soil was air-dried, sieved (<2 mm), and half of the material was sterilized twice by γ-irradiation at 18 kGy. Sterilized (S) and non-sterilized (NS) soils were stored at 4 °C until use. Soil was mixed with sand (0.71-1.25 mm) in a 1:1 ratio or 1:3 ratio (v/v) depending on the experiment.

### Plant material and growth conditions

*Arabidopsis thaliana* ecotype Columbia-0 (Col-0) seeds were used in all experiments. Seeds were surface-sterilized in 70% ethanol for 20 min, rinsed 5 times in sterile water, and stratified at 4 °C for 2-3 days. Seeds were sown directly onto pots containing the soil-sand mixtures described below. Plants were grown in a growth chamber (16 h light/8 h dark, 140 µmol m^-2^ s^-1^, 22 °C, and 55-60% relative humidity) for all experiments. Pots were watered from the bottom with deionized water as needed.

### Preparation of soil suspensions and dilution gradients

A soil suspension was prepared by mixing 20 g NS soil with 200 mL deionized water and shaking by hand for 1 min. This undiluted suspension was named D1 and was serially diluted 100-fold to generate 10^-2^ to 10^-10^ (D10^-2^-D10^-10^). This procedure was repeated three times. For a second experiment, a fresh D1 suspension was prepared by mixing 40 g NS soil with 400 mL water, shaken by hand and was passed through a 1 mm mesh to remove large particles and further diluted 2-, 4-, 8-, 12-, 16-, 20-, 24-, 32-, and 64-fold (D2-D64), with all suspensions sieved after each dilution. To rule out effects from dissolved nutrients, controls included a suspension prepared from sterilized soil (D1-S) and a suspension prepared from NS soil that was heat-treated at 80 °C for 1 h (D1-HT). For all inoculation experiments, 35 mL (for the D1-D10^-10^ series) or 40 mL (for the D1-D64 series) of each suspension was added to 400 mL S soil and mixed before combining with sand in a 1:3 ratio (v/v). Each soil preparation was distributed into five 9-cm pots and plants were grown for 5.5-6 weeks depending on the experiment.

### Experiment comparing S soil, NS soil and S soil inoculated with D2 suspension

*Arabidopsis* plants were grown in S soil, NS soil or in S soil inoculated with a D2 suspension as described above. For the S and NS treatments, deionized water (equivalent in volume to the added D2 suspension) was applied instead. For each treatment, nine 9-cm pots (each containing three plants) and three unplanted bulk-soil pots were prepared. The experiment was performed twice. Sampling of soil and plant material was conducted immediately after soil preparation (week 0) and after five weeks of plant growth. At week 0, bulk soil was collected from unplanted pots for nutrient measurements. At week 5, rhizosphere soil was obtained by recovering loosely adhering soil from the roots within each pot. Roots were pooled from each pot, rinsed with deionized water and blotted on sterile filter paper. Additional bulk soil was collected from unplanted pots. Rosette leaves and material for DNA extraction were snap-frozen in liquid nitrogen and stored at -80 °C before freeze-drying for 48-72 h. Bulk soil intended for nutrient analysis was stored at 4 °C until use.

### Leaf nutrient analysis

For the experiment comparing plants grown in S and NS soils, freeze-dried leaves were digested in 7% (v/v) HNO□, and analyzed for elemental concentrations by inductively coupled plasma optical emission spectrometry (ICP-OES). For the experiment comparing S, NS, and D2 treatments, whole rosettes were first homogenized in 15 mL tubes with 2-5 glass beads (3 mm) in a paint shaker for 2 min. Total nitrogen was quantified from 20 mg of ground rosette leaf tissue by the Dumas combustion method, and the remainder of the material was digested in 7% (v/v) HNO□ before ICP-OES.

### Soil nutrient analysis

Soil nitrate N (NO□^-^_-_N) and ammonium N (NH□□-N) were extracted from 4 g fresh soil with 16 mL 1 M KCl and quantified by flow-injection analysis (FIA). Inorganic P (Olsen P) was extracted from 2 g air-dried soil using 40 mL 0.5 M NaHCO□ (pH 8.5), acidified 1:5 (v/v) with 1.5 M H□SO□, and quantified by FIA. Exchangeable cations (Mg, Ca, K) were extracted from 2 g air-dried soil with 20 mL 1 M ammonium acetate (pH 7.0), acidified 1:1 (v/v) with 7% HNO□, and analyzed by ICP-OES.

### Nutrient supplementation experiment

Sterilized soil was mixed with sand in a 1:3 (v/v) ratio, and 340 g of the mixture was distributed into six 9-cm pots per tray. Each tray represented a single treatment. Nitrate was applied at 0, 60, 120, and 240 mg N kg^-1^ soil(Konishi et al. 2017), supplied as KNO_3_. Potassium concentrations were adjusted to the same level across treatments using KCl. In the control treatment (0 mg N kg^-1^ soil), K was supplied exclusively as KCl. For testing the effect of magnesium and calcium, nutrient solutions were prepared with 0.5 mM Ca supplied as CaCl_□_ ·2H_□_O, 0.25 mM Mg supplied as MgCl_□_ ·6H_□_O(Lemaître et al. 2008), a combination of both, or deionized water as control. Nutrient solutions were applied twice per week for 3 weeks (1 L per tray), starting 4 days after sowing. Rosette leaves were weighed after five weeks.

### Inoculation of organic agricultural soil with suspensions derived from alder swamp soil

Twofold (D2), fivefold (D5) and tenfold (D10) suspensions were prepared from the alder swamp soil using deionized water. For each treatment, 60 mL of each suspension was added to 600 mL of either autoclaved or non-autoclaved organic agricultural soil collected from Taastrup farm (Denmark) and mixed thoroughly. Deionized water was used as the mock treatment. Sand was then added at a 1:3 (soil:sand v/v) ratio. The resulting soil-sand mixtures were distributed into four to six 9-cm pots prior to sowing seeds.

### DNA extraction and sequencing

Freeze-dried root samples were ground with two stainless steel beads for 90 s at 30 s^-1^ using a TissueLyser II (Qiagen). DNA was then extracted from ground root material and freeze-dried soil using the DNeasy PowerSoil Pro Kit (Qiagen), following the manufacturer’s instructions. DNA quality and quantity were assessed using a NanoDrop spectrophotometer (Thermo Fisher Scientific, Waltham, MA, USA) and a Qubit fluorometer (Thermo Fisher Scientific), respectively. Library preparation and sequencing were performed by mBioWorks (Denmark).

For bacterial 16S rRNA gene amplicon sequencing, the V5-V7 region was amplified using the primers 799F (5′-AACMGGATTAGATACCCKG-3′) and 1193R (5′-ACGTCATCCCCACCTTCC-3′). Both the forward and reverse primers were tailed with sample-specific llumina index sequences to allow for deep NGS sequencing. The PCR was performed in a reaction mixture of DNA template 5-50 ng, 0.3μL forward primer (10μM), 0.3μL reverse primer (10μM), 5μLKOD FX Neo Buffer, 2μLdNTP (2 mM each), 0.2μL KOD FX Neo polymerase (Toyobo Life Sciences, Shanghai), and finally ddH2O added to a total volume of 20μL. After initial denaturation at 95 °C for 5 min, followed by 20 cycles of denaturation at 95 °C for 30 s, annealing at 50 °C for 30 s, and extension at 72 °C for 40 s, and a final step at 72 °C for 7 min. The amplified products were purified with an Omega DNA purification kit (Omega Inc., Norcross, GA, USA) and were quantified using Qsep-400 (BiOptic, Inc., New Taipei City, Taiwan). The insert sizes of sequencing libraries were inspected using a LabChip (PerkinElmer, USA). The amplicon library was paired-end sequenced (2×250) on an Illumina Novaseq 6000 sequencer.

For metagenomics, sequencing libraries were constructed using the NEBNext Ultra II DNA library Prep Kit for Illumina (New England Biolabs, Ipswich, USA). DNA was fragmented to an average size of 350 bp using a sonicator. The fragmented DNA was end-repaired, A-tailed, and ligated with Illumina-compatible adapters. The adaptor-ligated DNA was then amplified using PCR to enrich the library. The quality and size distribution of the libraries were assessed using an Agilent 2100 Bioanalyzer and quantification was performed using real-time PCR. The prepared libraries were sequenced on an Illumina NovaSeq 6000 platform (Illumina, San Diego, CA, USA) using a paired-end 150 bp (PE150) configuration, with a targeted sequencing depth of 10 Gb per sample.

### 16S amplicon sequencing analysis

The quality of the paired-end Illumina reads was assessed using FastQC v0.12.1(Andrews, Simon 2010). Primers were removed usingCutadapt v3.5(Martin 2011) with options ‘-e 0.1, -- overlap 8’. The reads were subsequently processed using USEARCH v11(Edgar 2010) as follows: low-quality reads were filtered to retain sequences with an error rate <1% (‘-fastq_filter’); redundant reads were removed using ‘-fastx_uniques’ with options ‘-minuniquesize 8, -sizeout’; and amplicon sequence variants (ASVs) were inferred using UNOISE3. Chimeric sequences were identified and removed by aligning ASVs to the SILVA (v138.2) database using the ‘-uchime2_ref’ command with options ‘-strand plus’ and ‘-mode balanced’. Host-derived sequences were filtered using the SINTAX algorithm with options ‘-sintax_cutoff 0.8’ and ‘-strand both’(Quast et al. 2013). After removing sequences annotated as mitochondria, chloroplasts, and eukaryotes, a total of 9,260 ASVs were retained. An ASV table was generated using USEARCH (‘-otutab’) with a minimum identity threshold of 97%, and taxonomic classification of ASVs was performed using the Ribosomal Database Project (RDP) classifier implemented in USEARCH (‘-sintax’)(Wang et al. 2007). The minimum sequencing depth across samples was 17,698 reads (Table S2), and the ASV table was rarefied to 17,000 reads per sample. Alpha and beta diversity analyses were conducted using USEARCH.

### Metagenomics analysis

For 134 metagenomic samples, *Arabidopsis* reads were removed using KneadData (v0.12.3; https://github.com/biobakery/kneaddata) with the options ‘SLIDINGWINDOW:4:20’ and ‘MINLEN:50’, in combination with Bowtie2 (‘--very-sensitive’) and the reference genome GCF_000001735.4_TAIR10.1 downloaded from NCBI(Sayers et al. 2025). Residual host-derived reads were further filtered by taxonomic classification using Kraken2 (v2.1.3) with the default plant database(Wood et al. 2019). Reads assigned to taxon identifier (taxid) 3702 (family Brassicaceae), including family-, genus- and species-level classifications, were removed. Cleaned samples with data sizes exceeding 6□GB were normalized to 6□GB. The normalized reads were assembled into 38,637,744 contigs using MEGAHIT (v1.2.9)(Li et al. 2015) with options ‘--min-contig-len 500’, ‘--k-min 37’, ‘--k-max 127’ and ‘--k-step 10’. Protein-coding genes were predicted using Prodigal (v2.6.3) (Hyatt et al. 2010), and redundant sequences were clustered and removed using MMseqs2 (Steinegger and Söding 2017) with parameters ‘--min-seq-id 0.95’, ‘-c 0.9’, ‘--cov-mode 1’ and ‘--cluster-mode 2’. The resulting non-redundant (NR) gene set was taxonomically classified using Kraken2 with a custom database comprising bacteria, archaea, fungi, viruses and UniVec_Coresequences. Paired-end clean reads from each sample were aligned to the NR gene set using Bowtie2, and high-quality alignments (identity ≥95% and alignment length ≥100□bp) were retained, converted to sorted BAM files, and used to calculate coding sequence (CDS) read counts with HTSeq-count (v2.0.9)(Anders et al. 2015). The NR gene sequences were translated into protein sequences and functionally annotated against the Kyoto Encyclopedia of Genes and Genomes (KEGG) database(Kanehisa et al. 2017) (v.102, April 2021 release) using DIAMOND BLASTP (v2.1.13.167)(Buchfink et al. 2015) with options ‘--outfmt 6’, ‘--max-target-seqs 1’, ‘--evalue1e-5’, ‘--sensitive’, ‘--block-size 4’ and ‘--index-chunks 1’. The resulting taxonomic and functional annotations of NR genes were further summarized to assess enrichment patterns (Table S6).

### Statistical analysis

Comparisons between two groups were performed using a Wilcoxon rank-sum test. Significance levels were indicated using asterisks (*, P < 0.05; **, P < 0.01; ***, P < 0.001). Differential abundance of KOs was assessed using the Wilcoxon rank-sum test, with P values adjusted for multiple comparisons using the Benjamini-Hochberg (BH) method. For comparisons involving multiple groups, a Kruskal-Wallis rank sum test was used, followed by Dunn’s post hoc test for pairwise comparisons. P values from multiple comparisons were adjusted using the BH method, with statistical significance defined as an adjusted P value < 0.05. For multiple-group comparisons, different letters were assigned based on adjusted P values to indicate significant differences between groups. All analyses were performed in R (v. 4.5.2) within RStudio (v. 2025.9.2). Data manipulation and visualization were conducted using the tidyverse package (v. 2.0.0), with figures generated using ggplot2 (v. 4.0.1) Statistical analyses were carried out using FSA (v. 0.10.0), rstatix (v. 0.7.3), and with multcompView (v.0.1.10) for Dunn’s post hoc tests.

## Author contributions

D.X. and Y.B. supervised the project; D.X. and L.D. conceptualized the experiments; L.D. and J.X. performed the experiments; X.Y conducted the 16S amplicon and metagenomics analyses; L.D. and D.X. analyzed the data and wrote the original draft; D.X., edited and reviewed the manuscript.

## Acknowledgements

We thank Camilla Timmermann Krogh for laboratory assistance, Elena Pupi, Ydun Kalsbeek-Hansen, Marlene Niedermayer for discussion of the data, the technical staff in the Section for Plant and Soil Science and the plant growth facilities at the Department of Plant and Environmental Sciences, University of Copenhagen, for their technical assistance. We acknowledge financial support from the Novo Nordisk foundation (NNF20OC0060824 to Barbara Ann Halkier).

## Data availability

The source data of each figure, the R-code for statistical analysis and plot generation are provided in the source data in R Markdown and HTML format.

## Supplementary Data

### Supplementary Figures

**Fig. S1** Soil microbiota alters shoot elemental composition in *Arabidopsis*.

**Fig. S2** Comparison of leaf elemental concentrations in non-sterilized and sterilized soil.

**Fig. S3** Serial dilution of soil microbiota identifies a growth-promoting community and rescues chlorosis.

**Fig. S4** Nitrate supplementation rescues plant growth and alleviates chlorosis.

**Fig. S5** Shannon evenness across soil and plant compartments.

**Fig. S6** Nitrogen cycling potential varies across soil and plant compartments.

**Fig. S7** D2 promotes plant growth in autoclaved organic agricultural soil.

### Supplementary Tables

**Table S1**. Soil nutrient concentrations in non-sterilized (NS), sterilized (S), and D2-inoculated soils.

**Table S2**. Metadata for soil, rhizosphere, and root samples used for 16S rRNA gene amplicon sequencing and metagenomics.

**Table S3**. PERMANOVA results based on Bray-Curtis dissimilarity matrices.

**Table S4**. Taxonomic profile at the phylum level in soil, rhizosphere, and root compartments based on 16S rRNA gene amplicon sequencing.

**Table S5**. Genes involved in inorganic nitrogen cycling and the degradation and mineralization of organic nitrogen.

**Table S6**. Metagenomic analyses for soil, rhizosphere, and root compartments.

